# Maternal glucocorticoid levels during incubation predict breeding success, but not reproductive investment, in a free-ranging bird

**DOI:** 10.1101/685891

**Authors:** Devin Fischer, Robby R. Marrotte, Eunice H. Chin, Smolly Coulson, Gary Burness

## Abstract

The hormone corticosterone (CORT)) has been hypothesized to be linked with fitness, but the directionality of the relationship is unclear. The “CORT-fitness hypothesis” proposes that high levels of CORT arise from challenging environmental conditions, resulting in lower reproductive success (a negative relationship). In contrast, the “CORT-adaptation hypothesis” suggests that, during energetically demanding periods, CORT will mediate physiological or behavioural changes that result in increased reproductive investment and success (a positive relationship). During two breeding seasons, we experimentally manipulated circulating CORT levels in female tree swallows (*Tachycineta bicolor*) prior to egg laying, and measured subsequent reproductive effort, breeding success, and maternal survival. When females were recaptured during egg incubation and again during the nestling stage, the CORT levels were similar among individuals in each treatment group, and maternal treatment had no effect on indices of fitness. By considering variation among females, we found support for the “CORT-adaptation hypothesis”; there was a significant positive relationship between CORT levels during incubation and hatching and fledging success. During the nestling stage CORT levels were unrelated to any measure of investment or success. Within the environmental context of our study, relationships between maternal glucocorticoid levels and indices fitness vary across reproductive stages.

**SUMMARY STATEMENT:** Maternal corticosterone levels predict breeding success of female tree swallows.

## INTRODUCTION

Within and among species individuals vary in the strategies used to maximise fitness, by adjusting the relative effort put into current versus future reproductive events (Williams, 2005; Hansen et al., 2016). There is ample evidence that increased energy expenditure and effort during one breeding bout results in decreased reproductive success, probability of re-nesting, or survival in subsequent bouts (Nager, 2006; Crossin et al., 2013, 2016; Harms et al., 2014; Bleu, Gamelon & Sæther, 2016; Henderson et al., 2017).

Glucocorticoids (GCs) have been hypothesized to be a mediator of the trade-off between current and future reproduction (Wingfield et al., 1998; Bleu, Gamelon & Sæther, 2016; Hansen et al., 2016). GCs are metabolic hormones that fluctuate daily with feeding and other activities, and under resting conditions regulate energy balance (Landys, Ramenofsky & Wingfield, 2006; Wilcoxen et al., 2011; Hau & Goymann, 2015). In response to an environmental stressor, GC levels increase rapidly, resulting in increased availability of metabolic substrates, and adjustment of behaviors toward immediate survival (Wingfield & Sapolsky, 2003; Romero, 2004) while inhibiting reproductive behaviour and physiology (Sapolsky et al., 2000; Dantzer et al., 2014), i.e. the CORT-trade-off hypothesis (Patterson et al., 2014).

GCs are thought to play a role in translating environmental cues into adaptive physiological responses. In birds, the dominant GC is corticosterone (hereafter, CORT), and an elevation of baseline CORT levels may signal a poor quality environment or an individual in poor condition (Bonier et al., 2009b). Following this reasoning, Bonier et al (2009a) formulated “the CORT-fitness hypothesis,” which predicts that individuals with higher circulating CORT levels would have lower fitness. In support of this, higher baseline CORT levels have been negatively associated with habitat quality, body condition, hatching success, and fledging success across various species, e.g. blue tits *Cyanistes caeruleus* (Henderson et al., 2017); tree swallows *Tachycineta bicolor* (Patterson et al., 2011); barn swallows *Hirundo rustica* (Saino et al., 2005); Adelie penguins *Pygoscelis adeliae* (Thierry et al., 2013).

In contrast, a positive relationship is predicted by “the CORT-adaptation hypothesis.” This hypothesis suggests that because CORT can mediate the mobilization of fuels, causing changes in behaviour or physiology that can increase investment in reproduction, elevated CORT will lead to higher fitness during energetically demanding times (Wingfield and Sapolsky, 2003; Bonier et al., 2009a). Indeed, across a variety of species and life history strategies individuals with higher reproductive success have been reported to have higher CORT levels, e.g. eastern bluebirds *Sialia sialis* (Burtka et al., 2016); black-legged kittiwakes *Rissa tridactlya* (Chastel et al., 2005); petrels *Macronectes* spp (Crossin et al., 2013); western bluebirds *S. mexicana* (Kleist et al., 2018); mourning doves *Zenaida macroura* (Miller et al., 2009).

In fact, there may exist no consistent relationship between CORT and fitness, due to a variety of factors masking directionality (Madliger and Love, 2016a). For example, a lack of relationship could be due to different functions of CORT; when resources are plentiful, elevated CORT could stimulate energy mobilization and parental provisioning; however, CORT could also be elevated in parents experiencing stressors (Vitousek et al., 2014). Even within a breeding season, different stages can have differing parental energetic requirements presumably requiring different levels of GC-mediated energy mobilization (Humphreys et al., 2006; Nager, 2006; Tulp et al., 2009; Sakaluk et al., 2018).

Most studies that have explored relations between CORT and fitness have been correlative. Although such studies are certainly informative, e.g. (Bonier et al., 2009b), studies that manipulate CORT levels directly, and explore the resultant parental response are also needed. Using tree swallows as a model organism, we attempted to elevate maternal CORT experimentally prior to egg-laying, and quantify subsequent variation in maternal reproductive investment and reproductive success over two breeding seasons. Females were captured shortly before egg laying and each received a silastic implant containing either corticosterone (CORT) or left empty (Sham). An additional group of females we captured for the first time during early incubation but received no implant. Although we refer to these latter females as “controls,” we recognize they may represent a non-random sample and we interpret our results accordingly. We asked two primary questions: (1) How does maternal CORT influence reproductive investment and success? (2) Does the directionality of the relationship between maternal CORT and reproductive investment and success change between the incubation and nestling provisioning stages, given the increased energy expenditure and brood value during chick provisioning? If the CORT-fitness hypothesis were true, we expected to detect a negative relationship between maternal CORT and measures of reproductive investment and success. Conversely, if the CORT-adaptation hypothesis were true, we expected to see a positive relationship between maternal CORT and reproductive investment and success.

## MATERIALS AND METHODS

### Study location and species

All protocols were approved by Trent University Animal Care Committee, with a handling, banding and collection permit provided by Canadian Wildlife Service, Environment Canada. Our study took place during spring and summer 2015 and 2016, using tree swallows, a small, migratory, aerial insectivore, that breeds across central and northern North America (Winkler et al., 2011). They are cavity nesters that readily occupy artificial nest boxes, and both sexes begin nest building in late April to early May, with laying occurring through May and June. Most populations lay only one clutch of 5 or 6 eggs per season; the female then incubates the eggs for about 14 days. Chicks are fed by both parents and fledge at approximately 21 days post-hatch.

We had two field sites near Peterborough Ontario, Canada (University Nature Area: 44 ° 21□ N, 78 ° 17□ W; and Lakefield Township Sewage Lagoon: 44 ° 25□ N, 78 ° 15□ W). In 2015 and 2016, the Nature Area had 66 and 70 boxes, respectively; Sewage Lagoon had 50 and 52 nest boxes. The Nature Area consisted of open woodland with long grasses, shrubs, and scattered apple (*Malus pumila)*, buckthorn (*Rhamnus cathartica)*, red cedar (*Juniperus virginiana*), white cedar (*Thuja occidentalis)*, and dogwood (*Cornus florida)*. The immediate landscape around boxes at the Sewage Lagoon was exposed soil and grasses; the greater surrounding area was farmland consisting of both crop and pasture land. Nest boxes at the Sewage Lagoon were 5 to 10 metres from the water.

### Experimental manipulation of maternal corticosterone levels

Nest boxes were monitored daily beginning 6 May in both years. When nests were about 75% formed (when cup-shaped or when feathers were present), we captured females using cardboard trapdoors over the nest box opening, or by surprising birds sitting in nest boxes. In 2015, seven females were caught at night (between 2200 and 2400 hours) by surprising birds sitting in nest boxes (no females were found in nest boxes at night in 2016). Upon capture, females were randomly assigned to CORT or sham treatment groups (Table 1). We sterilized the skin of the right flank with 70% ethanol, made a 4mm subcutaneous incision, and inserted a sterilized 10mm silastic tube (ID 1.47mm and OD 1.96mm, Dow Corning 508-006) filled with crystalline CORT (Sigma Aldrich C2505) that was sealed with silicone sealant (732 Dow Corning) at both ends (CORT treatment). To each implant we added a single hole using a 30 G needle (Ouyang et al., 2013). The design of our implants followed that of Ouyang et al. (2013), who used a 7 mm long implant (ID 1.5 mm), sealed at both ends, and punctured with a single 0.3 mm hole. In great tits (*Parus major*) this design increased corticosterone levels by ∼2-fold above baseline for approximately 30 days post-implantation (Table 1 in Ouyang et al. 2013). Sham treatment tree swallows received sterilized empty implants. Empty implants weighed approximately 0.02g and held an average of 0.007g ± 0.0007g of CORT. Once the implant was inserted, the incision in the skin was sealed with a drop of 3M Vetbond (no. 1469SB). Each female was then aged as second year, SY, or after-second year, ASY (Pyle et al., 1987). Flattened wing length was measured with a standard ruler with a wing stop (±1mm), mass was measured with a Pesola spring scale (±0.25g). All birds (including any males caught inadvertently) were banded with a federal aluminum numbered leg band (Canadian Wildlife Service) and released. Birds were held for no more than 10 minutes before release. In 2016 and 2017, any previously banded female from 2015 or 2016 was counted as a returned bird in the return rate analysis regardless of whether they hatched a clutch that year.

**Table 1.** Sample sizes of adult female tree swallows allocated to each maternal treatment group across two years.

|  | Sham |  | CORT |  | Control |  |
| --- | --- | --- | --- | --- | --- | --- |
|  | SY | ASY | SY | ASY | SY | ASY |
| Number of females implanted | 16 | 27 | 18 | 26 | - | - |
| Number of females that laid | 3 (9) | 17 (24) | 5 (6) | 9 (18) | 11 | 10 |
| Number of females that laid eggs that hatched | 2 (7) | 16 (23) | 4 (5) | 9 (16) | 8 | 10 |

Sham: females had an empty silastic implant; CORT: females had a silastic implant filled with crystalline corticosterone; Control: females had no implant.
Female age: SY (second-year); ASY (after second-year).
Bracketed values represent total number of individuals handled/implanted; non-bracketed values indicate sample sizes of birds with implants that were still present when the bird was recaptured during incubation.
Four females of unknown age are omitted from the table (1 CORT, 1 Sham, 2 Controls)

We allocated females to the Control group if they were not caught prior to laying, either because they did not enter the nest box while it had a trap, or because they began laying earlier than we expected. Although these females did not receive an implant before egg laying they were handled and measured beginning during incubation (sample sizes in Table 1).

### Nest and egg monitoring

Nest boxes were monitored daily throughout the nest-building and laying period, and when eggs were discovered, eggs were numbered with a black marker and weighed (±0.01 g) using a digital balance. One female had a lay date of 15 June, which was greater than 3 standard deviations from the population mean (21 May). We did not consider this female further because we suspected it was re-nesting after a failed first attempt. All other nests were included in statistical analyses.

### Nestling measurements

Beginning on day 12 of incubation (incubation day 0 = first day no new eggs were laid, and eggs were warm to the touch), nest boxes were checked twice daily. The hatch day of the first nestling was defined as day 0 for that nest. It was not possible to match nestlings to egg identity. We marked the talons of nestlings with coloured nail polish to distinguish individuals, until we banded them on day 10 post-hatch with aluminum numbered leg bands (Canadian Wildlife Service). Nestlings were weighed at hatch with an egg scale (± 0.01g), and on days 3, 7, 10, 13, and 14 post-hatch with a Pesola spring scale (±0.25g). Beginning on day 18 post-hatch, we checked nest boxes daily by partially opening the door to determine fledging success. To guard against pre-mature fledging, the nest box opening was blocked for 1 minute after checking, and when the blocking was removed the box was observed for 5 min from a distance of a few metres; no instances of premature fledging were observed.

### Blood sampling procedure

We recaptured adult females in nest boxes between day 2 and 5 of incubation (both years) and between day 3 and 6 post-hatch during chick rearing (in 2016 only) between 0600 and 1200 hours. Upon capture, we collected a 100µl blood sample from the brachial vein using a micro- capillary tube within three minutes of the female entering the nest box. The mean time taken to draw blood (±SE) was 125 ± 5s (N=61) during incubation, and 115 ± 6s (N=26) during the nestling stage. Blood samples were kept on ice for up to eight hours. Samples were then centrifuged for four minutes at 19,200 x *g* (Thermo IEC Micro-MB) before plasma and red blood cells were frozen separately at −80°C. Prior to release, we recorded female body mass and marked the tail feathers and right primaries with a spot of white acrylic paint to distinguish females from males during subsequent behavioural observations (Whittingham et al., 2003; Bonier et al., 2011). If the female had not been captured previously (i.e., she was to become a Control female), upon first capture during incubation we recorded her head-bill length and wing length, and banded her.

We collected nestling blood samples (50µl) from the brachial vein on days 7 or 8, and 13 post-hatch. Samples taken on day 7 or 8 post-hatch were for molecular sexing and were added to 1ml of lysis buffer in the field and subsequently stored at −20°C. Samples collected on day 13 were centrifuged and plasma was stored at −80°C (as part of a separate study).

### Adult behavioural observation

On day 7 or 8 post-hatch between 0830 and 1400 hours, nest boxes were observed from a distance of 10 m for 1 hr (Lendvai et al., 2015), during which we counted the number of visits made by males and females to the nest box. This was the maximum distance at which it was still possible to distinguish the sex of the adult entering the box through binoculars. Observations made mid-day have been shown to provide the best estimates of feeding rate, although 1-hour observations periods done at any time of day predict total daily feeding rates (Lendvai et al., 2015)

### Lab procedures

#### Corticosterone radioimmunoassay

Plasma samples were analyzed for total corticosterone in duplicate using a ^125I^ radioimmunoassay (MP Biomedicals #07120103) following the manufacturer’s instructions (Washburn et al., 2002). This assay has low cross-reactivity with deoxycorticosterone (0.34 %), testosterone (0.10 %), cortisol (0.05 %), aldosterone (0.03 %), and progesterone (0.02%). Plasma was diluted 1:25 (10µl of plasma plus 240µl of assay buffer). Samples that were not detectable were set to the lowest point on the standard curve (3.125 ng/ml), following Hogle and Burness (2014). We did not extract plasma because a serial dilution of non-extracted plasma pooled from five individuals was parallel to the standard curve. A total of 23 individual assays was performed. To calculate the inter-assay coefficient of variation (CV) we ran duplicates of the kit “low” and “high” controls in each assay. The inter-assay CV was 8.6% and 7.4% for the low and high controls, respectively. To calculate the intra-assay CV, in a single assay we included 4- replicates of the low and high controls. The intra-assay CVs were 13.4% and 7.9% (respectively).

#### Molecular sexing protocol

Maternal CORT may facilitate sex-biased investment in nestlings (Love et al., 2005; Love and Williams, 2008). To evaluate this possibility, nestling blood samples taken on day 7 or 8 post-hatch were used for genetic sexing using the CHD1W and CHD1Z genes (Fridolfsson and Ellegren, 1999; Hogle and Burness, 2014). DNA extraction was done using DNEasy blood and tissue kits (Qiagen 69506). A touchdown PCR procedure was used with 10ul volumes consisting of 1.2ul 10X buffer, 0.4ul MgCl_2_, 1.0ul dNTP, 0.25ul BSA, 0.2µl each of primers 2550 and 2718 (Fridolfsson and Ellegren, 1999), 1.0µl Taq polymerase, 3.75µl H_2_O, and 2µl DNA in an Eppendorf thermocycler. Initial denaturing began at 94°C for 5 min followed by a touchdown sequence where the annealing temperature was lowered 1°C per cycle from 94° to 50°C. A further 24 cycles were run with a denaturing temperature of 94°C for 30 s, annealing temperature of 40°C for 30 s and extension of 72°C for 30 s, followed by a final extension at 72°C for 2 min after the last cycle. PCR products were separated in a 3% agarose gel stained with ethidium bromide and run in 1X TBE buffer. Each gel was run with known male and female adult samples for comparison (N=273 chicks).

### Statistical analyses

All data will be deposited in the DataDryad data repository. We used R version 3.4.3 (2017) to run all analyses, and statistical significance was claimed at P < 0.05. During field work we were generally blind to the experimental treatment, but not during statistical analysis. Sample sizes were determined by the number of breeding individuals in our study population that could be captured. To improve normality, all CORT values were log_e_ transformed; all other metrics were untransformed. Raw means are reported ± SE. Sample sizes varied among analyses because we were not always able to collect all measurements from all individuals. We included ‘year’ as a factor only in analyses of maternal CORT during incubation (CORTinc), because during the nesting phase (CORTnest) we measured CORT in one year only (2016). To avoid possible carry- over effects of experimental treatment, females re-captured during the second year of the study were included for their first year of capture only.

We constructed our statistical models including only main effects that were of likely biological importance and/or *a priori* interest; as such, not all two-way interactions were included. We report outputs from global statistical models. Because we had explicit hypotheses, and because none of our response variables was correlated, we did not to use a post-hoc correction for the number of tests performed (Perneger, 1998; Streiner, 2015).

#### Morphological and hormonal measures of adult females

We ran preliminary tests to determine whether females that had been assigned to CORT or Sham treatment groups differed in pre-implant body mass (measured at time of implant; females in the Control group were not captured prior to incubation and thus there was no pre-implant mass measurement). To test for possible differences in body size among treatments, we compared a female’s wing length (measured pre-laying in the CORT and Sham treatments, and during early incubation in the Control females). Finally, we tested for differences in clutch initiation date in Julian days among treatments. Separate linear models (LM) were run with female pre-laying mass, wing length, and clutch initiation date as the response variable, and treatment (CORT, Sham, Control), site (Nature areas, Sewage Lagoon), year (2015, 2016), and age (Second year, SY; After second year, ASY) as the predictors. We did not include any interactions terms as they were not of *a priori* interest.

To test whether implanted females differed in their probability of recapture depending on treatment or year, we ran a generalized linear model (GLM) with binomial errors, with recapture status (recaptured/non-recaptured) as the dependent variable, and treatment and year as the fixed effects. To test whether the total number of individuals that retained their implants and subsequently laid eggs differed between the CORT and Sham maternal treatment groups, we used a chi-square test (because Control females were only captured post-egg laying, they were not included in this analysis).

#### Maternal baseline corticosterone during incubation and nestling stages

To test whether treatment affected maternal CORT levels within each breeding stage (incubation and nestling), we used linear models (LM) with either CORT during incubation (from hereafter CORTinc) or CORT during the nestling stage (CORTnest) as the response variable and maternal treatment, age, site, sample time (time from initial contact with bird to end of blood sample), and clutch initiation date (in Julian days) as fixed effects. We had no *a priori* predictions regarding interactions, so none was included in the models.

We analyzed CORTinc and CORTnest separately because CORTnest was only measured in 2016. Baseline CORTinc measurements (N=59) had one suspected outlier (121.22 ng/ml) removed prior to analysis. This value was > 3 standard deviations from the mean; considerably higher than the 0.5 to 14 ng/ml range reported for previously (Franceschini et al., 2008; Ouyang et al., 2011; Patterson et al., 2011; Madliger et al., 2015). Preliminary analyses were run with and without this outlier, and although no difference was found in the pattern of significance of parameters, we chose to exclude it.

#### Measures of female reproductive investment

As indices of maternal investment during incubation we used clutch mass (summed mass of individual eggs at laying), and during the nestling phase we used maternal nest box visitation rate and nestling growth rate. To test whether a female’s clutch mass correlated with her CORT levels, we used a linear model (LM) with clutch mass as the response variable and CORTinc, maternal treatment, age, site, and year as main effects. To explore investment during the nestling stage, we used a LM with the number of nest box visits per chick per hour (by the female) as the response variable and CORTnest, treatment, maternal age, site, and male nest box visits per chick per hour as fixed effects. Finally, we calculated nestling growth rate per day during the linear growth phase (Burness et al., 2001) as the difference in individual mass between days 3 and 7 post-hatch, divided by 4 days. We used a linear mixed model (LMM, lmer in R package lme4) with individual chick mass gain per day as the response variable, and nest ID as a random effect. Fixed effects were CORTnest, maternal treatment, maternal age, site, nestling sex. To evaluate the possibility that maternal corticosterone may be linked with sex-specific investment in offspring (e.g., Love et al., 2005), we included an interaction between nestling sex and CORTnest.

#### Measures of female reproductive success

To test for a relationship between CORTinc and indices of reproductive success, we used a generalized linear mixed model (GLMM; glmer in R package lme4) with binomial errors, with either hatching or fledging success as the response variable (0 or 1 for each chick) and CORTinc, maternal treatment, age, site, and year as fixed effects, and nest ID as a random effect. To explore the relationship between CORTnest and post-hatching reproductive success, we examined individual nestling mass at day 14 post-hatch and fledging success as indices of reproductive success. To test whether nestling mass differed with maternal CORT or treatment, we used a LMM with nestling mass at day 14 as the response variable and CORTnest, maternal treatment, maternal age, and site as fixed effects (year was not included because CORTnest was measured in 2016 only), and Nest ID as a random effect. Finally, to test whether fledging success differed with maternal CORT or treatment, fledging success (0 or 1 for each chick) was used as the response variable in a GLMM with binomial errors with maternal treatment, maternal age, site, and CORTnest as fixed effects, and Nest ID as a random effect. No interaction terms were included in these analyses.

#### Measures of female survival

We estimated female survival by using the return rates of adult females to the study sites the following spring and comparing this with CORTinc or CORTnest during the previous year in separate models. Return rate (either 0 or 1) was the response variable in a general linear model (GLM), with CORTinc (or CORTnest), treatment, year, age, site, and number of nestlings fledged as main effects. In analyses of CORTnest, “year” was not included in the model because CORTnest was only measured in a single year (2016).

## RESULTS

### Morphology and hormonal measures of adult females

We implanted 45 females with corticosterone-filled implants (CORT), and 44 with sham implants (Sham); an additional 23 females were captured for the first time during incubation and were allocated to the Control treatment (Table 1). There was no difference in pre-egg laying body mass between females allocated to the CORT and Sham groups (Table S1); females in the Control group were not captured prior to egg laying, so there was no pre-egg laying mass. Focusing on individuals that retained their implants, wing length and clutch initiation date did not differ significantly among treatments (Table S1). There was no significant difference between the Sham and CORT treatments in the percentage of females that retained their implants and subsequently laid eggs (Sham: 45% (20 of 43), CORT: 31% (14 of 44); χ^2^=0.865, df=1, p = 0.352; Table 1; a single CORT and Sham female of unknown age were omitted from the analysis).

There was no significant difference by year in the number of implanted females that were recaptured during incubation (β = 0.800, SE = 0.509, z = 1.573, p = 0.116, N_2015_=34 recaptured and 25 non-recaptured, N_2016_=22 recaptured and 8 non-recaptured. However, sham-implanted individuals were more likely to be recaptured than CORT-implanted (β = 1.115, SE = 0.468, z = 2.381, p = 0.017, N_Sham_=33 recaptured and 11 non-recaptured, N_CORT_=23 recaptured and 22 non-recaptured). Control birds were not included in the recaptured/not recaptured analysis because they were caught for the first time during incubation.

### Implants failed to raise long-term maternal corticosterone levels

During incubation, females were recaptured on average 17.02 days (±0.63) after implantation (range 7 to 26 days). Contrary to expectations, when females were recaptured there was no difference in CORT levels among the 3 treatments (Table 2; Fig. 1A). Lay date (i.e., clutch initiation date) was also not a significant predictor of CORTinc (Table 2). However, older mothers (ASY) had higher CORTinc levels than SY mothers and levels differed between years (Table 2). During nestling provisioning, maternal baseline CORT (CORTnest) did not differ among treatments (Table 2, Fig. 1B), nor with any other fixed effects (Table 2).

**Table 2.** Factors contributing to variation in corticosterone levels in female tree swallows during incubation (CORTinc) and the nestling stage (CORTnest).

| Response variable | Fixed effects | $\beta$ | SE | df | t | P | R<sup>2</sup> |
| --- | --- | --- | --- | --- | --- | --- | --- |
| CORTinc<br>(ng/ml) | Intercept | 2.753 | 3.578 | 1, 38 | 0.769 | 0.447 | 0.227 |
|  | Treatment (CORT) | -0.611 | 0.315 | 1, 38 | -1.943 | 0.060 |  |
|  | Treatment (Sham) | -0.366 | 0.276 | 1, 38 | -1.325 | 0.193 |  |
|  | Age (ASY) | 0.770 | 0.327 | 1, 38 | 2.358 | **0.024** |  |
|  | Lay date | 0.000 | 0.025 | 1, 38 | 0.008 | 0.993 |  |
|  | Site (Nature Area) | -0.459 | 0.255 | 1, 38 | -1.800 | 0.080 |  |
|  | Sample time | -0.005 | 0.004 | 1, 38 | -1.233 | 0.225 |  |
|  | Year | -0.755 | 0.291 | 1, 38 | -2.591 | **0.014** |  |
| CORTnest<br>(ng/ml) | Intercept | 2.834 | 4.272 | 1, 17 | 0.664 | 0.516 | 0.071 |
|  | Treatment (CORT) | 0.593 | 0.369 | 1, 17 | 1.610 | 0.126 |  |
|  | Treatment (Sham) | 0.218 | 0.292 | 1, 17 | 0.748 | 0.465 |  |
|  | Age (ASY) | 0.086 | 0.489 | 1, 17 | 0.176 | 0.862 |  |
|  | Lay date | -0.005 | 0.028 | 1, 17 | -0.163 | 0.872 |  |
|  | Site (Nature Area) | 0.336 | 0.362 | 1, 17 | 0.926 | 0.367 |  |
|  | Sample time | 0.002 | 0.004 | 1, 17 | 0.638 | 0.532 |  |

Year was not included for CORTnest, because data were collected in a single year. Statistically significant main effects are in bold.

**Figure 1.**
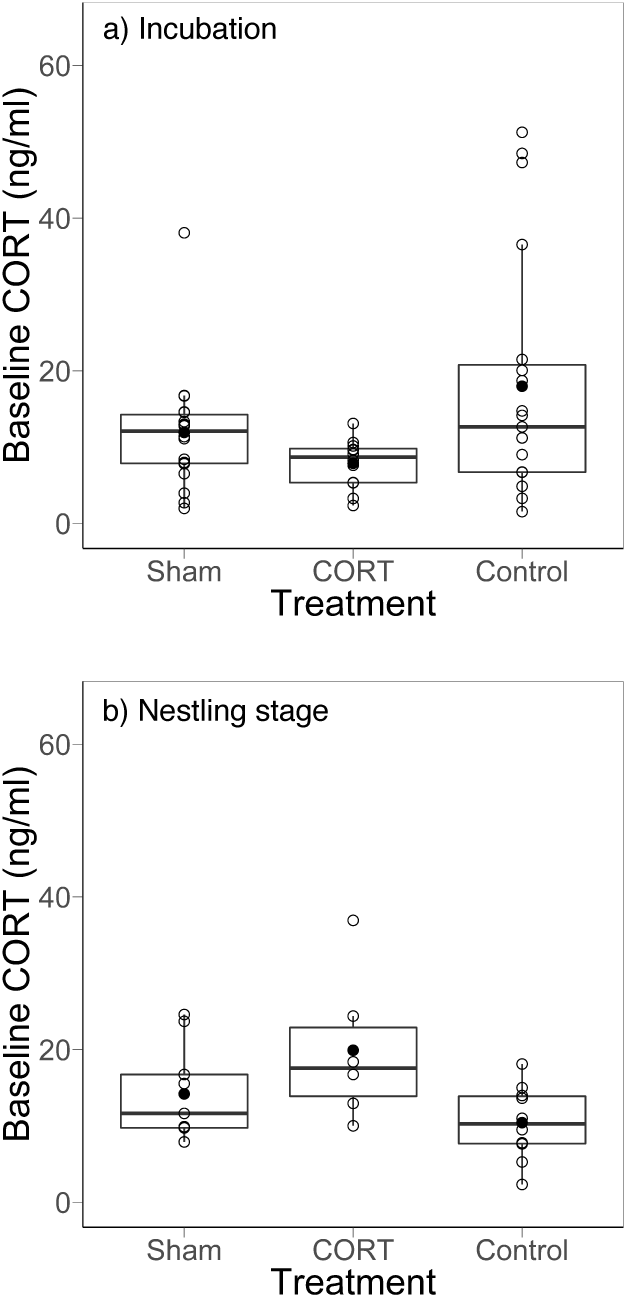
There was no significant difference among treatments in plasma corticosterone levels of female tree swallows when measured during (A) incubation and (B) nestling stage. Individuals in the CORT treatment had a single silastic implant containing crystalline corticosterone, those in the Sham treatment had an empty implant, while Control birds had no implant. The black circle indicates the mean; the thick horizontal line is the median. Individual data points are shown.

### Maternal corticosterone levels did not predict reproductive investment

Mean clutch size (±SE) of females was 5.3 eggs ± 0.1 (range = 3 to 7 eggs per nest, N=67 nests). Reproductive investment during laying, measured as clutch mass, did not correlate with maternal CORT levels during incubation (CORTinc) nor with maternal treatment, although older birds had significantly heavier clutches (Table 3). Similarly, during the nestling stage, there was no relationship between either maternal CORT (CORTnest) or treatment on the number of female nest box visits (Table 3). Although maternal treatment did not influence nestling growth rate between days 3 and 7, there was a marginally significant negative relationship between maternal CORTnest and nestling growth rate (p = 0.092, Table 3). Maternal age influenced nestling growth rates, with nestlings from SY mothers having higher growth rates than nestlings from ASY mothers (Table 3).

**Table 3.**
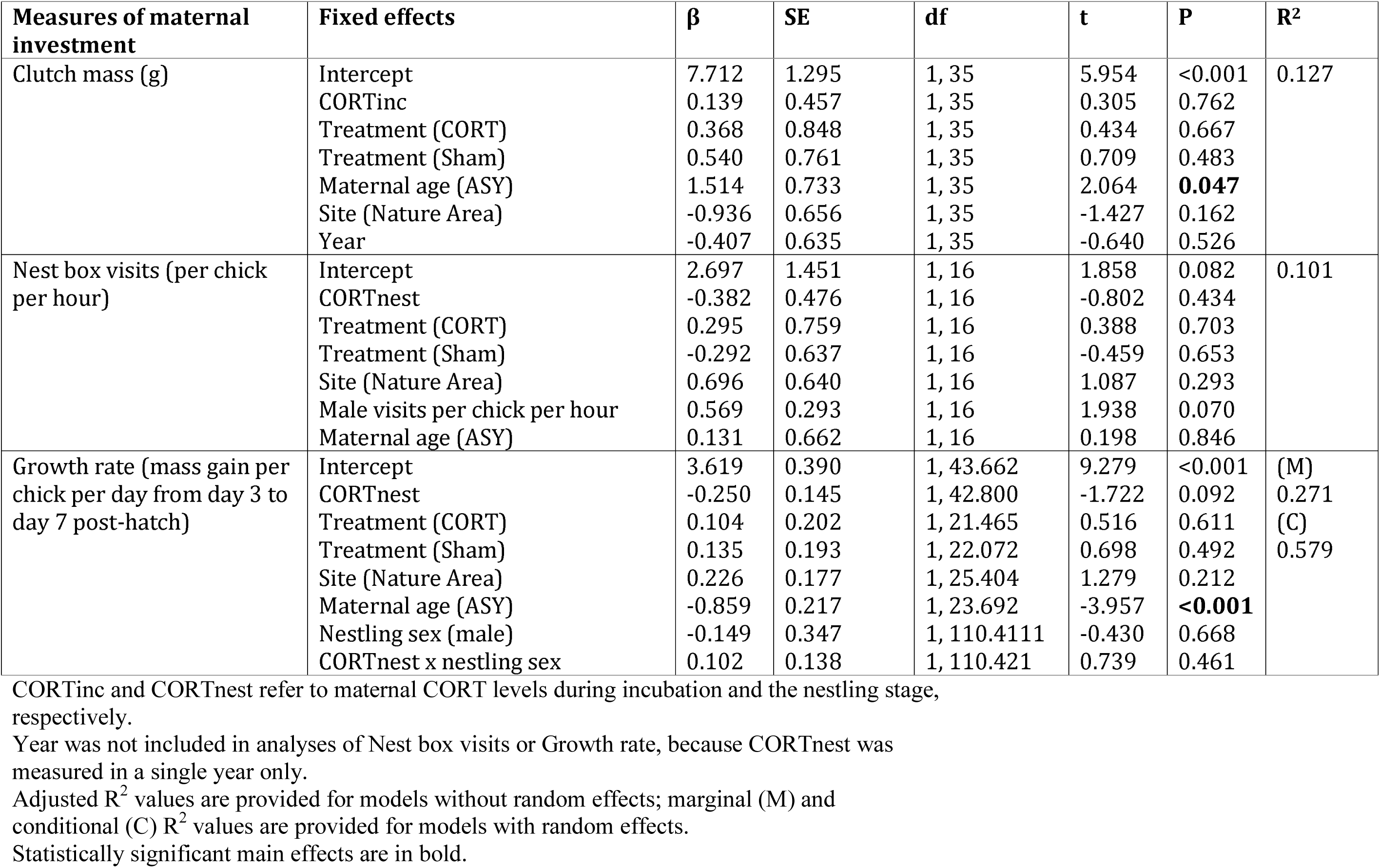
Factors contributing to variation in reproductive investment in female tree swallows.

### Maternal corticosterone levels during incubation predicted reproductive success

As indices of reproductive success, we measured hatching success, nestling mass at day 14 post-hatch, and fledging success. Mothers with higher CORT during incubation (CORTinc) had significantly higher hatching success (Table 4). Nestling mass at day 14 post-hatch was not predicted by either CORTnest nor maternal treatment, although nestlings at the Nature Area site tended to be heavier (Site: p = 0.064; Table 4).

**Table 4.** Factors contributing to variation in reproductive success in female tree swallows.

| Measures of maternal reproductive success | Fixed effects | $\beta$ | SE | df | z or t | P | R<sup>2</sup> |
| --- | --- | --- | --- | --- | --- | --- | --- |
| Hatching success | Intercept | -0.831 | 1.046 | 1, 263 | -0.795 | 0.427 | (M) 0.220<br>(C) 0.461 |
|  | CORTinc | 0.969 | 0.378 | 1, 263 | 2.563 | **0.010** |  |
|  | Treatment (CORT) | 1.260 | 0.744 | 1, 263 | 1.694 | 0.090 |  |
|  | Treatment (Sham) | 0.418 | 0.688 | 1, 263 | 0.609 | 0.543 |  |
|  | Site (Nature Area) | -0.818 | 0.572 | 1, 263 | -1.430 | 0.153 |  |
|  | Year | -0.285 | 0.530 | 1, 263 | -0.538 | 0.591 |  |
|  | Age (ASY) | 0.599 | 0.598 | 1, 263 | 1.002 | 0.316 |  |
| Nestling mass at day 14 post-hatch | Intercept | 22.219 | 0.963 | 1, 29.767 | 23.082 | <0.001 | (M) 0.086<br>(C) 0.239 |
|  | CORTnest | -0.587 | 0.363 | 1, 26.735 | -1.618 | 0.117 |  |
|  | Treatment (CORT) | 0.291 | 0.592 | 1, 22.059 | 0.492 | 0.628 |  |
|  | Treatment (Sham) | -0.356 | 0.575 | 1, 21.742 | -0.619 | 0.543 |  |
|  | Site (Nature Area) | 0.972 | 0.503 | 1, 27.232 | 1.934 | 0.064 |  |
|  | Age (ASY) | 0.226 | 0.600 | 1, 27.066 | 0.376 | 0.710 |  |
| Fledging success A | Intercept | -2.437 | 1.450 | 1, 228 | -1.681 | 0.093 | (M) 0.333<br>(C) 0.670 |
|  | CORTinc | 1.448 | 0.551 | 1, 228 | 2.627 | **0.009** |  |
|  | Treatment (CORT) | 0.916 | 0.998 | 1, 228 | 0.918 | 0.359 |  |
|  | Treatment (Sham) | 0.538 | 0.927 | 1, 228 | 0.580 | 0.562 |  |
|  | Site (Nature Area) | -1.542 | 0.826 | 1, 228 | -1.868 | 0.062 |  |
|  | Age (ASY) | 1.029 | 0.849 | 1, 228 | 1.212 | 0.225 |  |
|  | Year | -0.491 | 0.684 | 1, 228 | -0.718 | 0.473 |  |
| Fledging success B | Intercept | 6.423 | 2.922 | 1, 144 | 2.198 | 0.028 | (M) 0.186<br>(C) 0.591 |
|  | CORTnest | -2.029 | 1.078 | 1, 144 | -1.883 | 0.060 |  |
|  | Treatment (CORT) | 0.207 | 1.295 | 1, 144 | 0.160 | 0.873 |  |
|  | Treatment (Sham) | -0.653 | 1.190 | 1, 144 | -0.549 | 0.583 |  |
|  | Site (Nature Area) | -0.325 | 1.202 | 1, 144 | -0.270 | 0.787 |  |
|  | Age (ASY) | 0.819 | 1.461 | 1, 144 | 0.561 | 0.576 |  |

Each model includes Maternal ID as a random effect; marginal (M) and conditional (C) R<sup>2</sup> values are provided.
There were two analyses of Fledging success (A and B), with predictors including either **CORTinc** or **CORTnest**, respectively.
Year was not included in analyses of Nestling mass at day 14 or Fledging success B, because **CORTnest** was measured in a single year only.
Statistically significant main effects are in bold.

The probability of a nestling fledging significantly increased with maternal corticosterone levels measured during incubation (Fledging success A, Table 4). There was a marginally significant negative relationship during the nestling phase (Fledging success B, p = 0.060), but this was driven, at least in part, by a control female with the lowest hormone levels (2.34 ng/ml) yet 100% fledging success. Maternal treatment had no effect on fledging success (Fledging success A or B, Table 4). Fledging success tended to be higher at the Sewage Lagoon site (Fledging success (A), Site: p=0.062; Table 4).

### Maternal return rate was not significantly predicted by maternal corticosterone levels

Twenty-nine of 67 (43%) females (Sham, CORT or Control) returned in the year after they were initially caught, and all returning females returned to the same breeding site where initially caught. The number of females included in the analysis differs from totals in Table 1, because only females with corticosterone measurements were included. The probability that a female returned tended to increase with her incubation CORT levels (CORTinc, p = 0.054) and number of fledglings in the previous year (Return rate (A): p=0.056, Table 5).There was no significant effect of CORTnest or maternal treatment on the likelihood of a female returning to the nest sites the following year (Table 5).

**Table 5.** Factors predicting the return rate in female tree swallows in the following year.

| **Response variable** | **Fixed effects** | **$\beta$** | **SE** | **df** | **z** | **P** |
| --- | --- | --- | --- | --- | --- | --- |
| Return rate A | Intercept | -3.738 | 1.686 | 1, 34 | -2.217 | 0.027 |
|  | CORTinc | 1.177 | 0.612 | 1, 34 | 1.925 | 0.054 |
|  | Treatment (CORT) | 1.104 | 1.063 | 1, 34 | 1.038 | 0.299 |
|  | Treatment (Sham) | 1.049 | 0.962 | 1, 34 | 1.091 | 0.275 |
|  | Number of fledglings | 0.538 | 0.281 | 1, 34 | 1.913 | 0.056 |
|  | Year | 0.240 | 0.820 | 1, 34 | 0.293 | 0.769 |
|  | Age (ASY) | -2.077 | 1.069 | 1, 34 | -1.943 | 0.052 |
|  | Site (Nature Area) | 0.758 | 0.939 | 1, 34 | 0.808 | 0.419 |
| Return rate B | Intercept | -2.562 | 3.211 | 1, 15 | -0.798 | 0.425 |
|  | CORTnest | 0.337 | 1.066 | 1, 15 | 0.317 | 0.752 |
|  | Treatment (CORT) | 0.579 | 1.678 | 1, 15 | 0.345 | 0.730 |
|  | Treatment (Sham) | 0.570 | 1.262 | 1, 15 | 0.452 | 0.651 |
|  | Number of fledglings | 0.636 | 0.409 | 1, 15 | 1.557 | 0.119 |
|  | Age (ASY) | -0.730 | 1.545 | 1, 15 | -0.472 | 0.637 |
|  | Site (Nature Area) | 1.073 | 1.542 | 1, 15 | 0.696 | 0.487 |

There were two analyses of Return rate (A and B), with predictors including either CORTinc or CORTnest, respectively.
Year was not included in analyses of Return rate B, because CORTnest was measure in one year only.
Statistically significant main effects are in bold.

## DISCUSSION

Our data support the CORT-adaptation hypothesis. During egg incubation, corticosterone levels of female tree swallows were positively related to two measures of reproductive success (hatching and fledging success) and positively (albeit non-significantly) with female return rates. During the nestling stage, there was no relationship between corticosterone and indices of either reproductive investment or reproductive success. During neither period did we detect a significant negative relationship between CORT and fitness, as predicted by the CORT-fitness hypothesis.

### Maternal corticosterone levels during incubation

We tested for a relationship between maternal corticosterone levels during incubation (CORTinc) and clutch mass (as a single measure of reproductive investment), and hatching success and survival to fledging as measures or reproductive success (Bonier et al., 2009b; Schoenle et al., 2017). We found no relation between CORTinc and clutch mass, however, female tree swallows with higher CORTinc levels had greater hatching success and higher fledging success. The positive relationship we detected with hatching success may be due to CORT mobilizing energy stores, and thus allowing for increased reproductive effort (Riechert et al., 2014). However, positive (common terns *Sterna hirundo)* (Riechert et al., 2014), negative (zebra finches) (Khan et al., 2016), and null relationships (red-winged blackbirds) (Schoenle et al., 2017) have all been reported between maternal CORT and hatching success. Differences in the directionality of the relationships are presumably due to various environmental factors, including weather conditions (Schoenle et al., 2017), food (Riechert et al., 2014), and/or resource availability (Breuner and Burk, 2019).

A positive relationship between CORTinc and fledging success is consistent with relationships reported in eastern bluebirds (Burtka et al., 2016) and blue tits (Henderson et al., 2017). However in tree swallows, both negative (Bonier et al., 2009b) and statistically non-significant (Madliger and Love, 2016a) relationships between CORTinc and number of fledglings have been reported. A positive relationship, such as we detected, between maternal CORT during incubation and fledging success might be expected if the relationship were mediated through maternal transfer of CORT into the egg, leading to higher begging rates and body size in nestlings of mothers with higher CORT (Bowers et al., 2016). However, this would be a plausible mechanism only if CORT levels during incubation correlated with levels pre-laying, as has been found in other tree swallow populations (Ouyang et al., 2011); something we did not evaluate in our study.

A positive relationship between maternal CORT and fitness (CORT-adaptation hypothesis) should emerge when CORT levels are increased to meet higher energetic demands associated with reproduction (Bonier et al., 2009a; Crossin et al., 2013; Rivers et al., 2017). During incubation, individuals may experience more unpredictable stressors than during the nestling stage (Romero, 2002). For example, challenging environmental conditions such as lower temperatures and scarcer food resources in early spring can cause a negative relationship between both temperature and foraging success and baseline CORT levels, depending on the fitness and environmental measure used (Angelier et al., 2007; Wingfield et al., 2010; Ouyang et al., 2015). Because higher baseline levels may prime the body to perform better under stress, females with higher baseline CORT during incubation in our study may have been better able to meet these challenges (Romero, 2002).

### Maternal corticosterone levels during chick rearing

We predicted that if there were a relationship between CORT and reproductive investment and success, it would most likely emerge post-hatch, given the higher maternal energy expenditure required during chick rearing than during incubation (Nilsson and Råberg, 2001; Humphreys et al., 2006; Sakaluk et al., 2018) but see (Williams, 2018). However, female CORT levels during chick rearing were unrelated to any measure of reproductive investment (nest box visits and and nestling growth rate) nor any measure of reproductive success (nestling mass at day 14 and fledging success). Despite our inability to detect relationships, others have reported that individuals with higher baseline CORT levels during chick rearing had higher parental foraging effort, provisioning rates, and energy transfer to the nestlings, e.g. macaroni penguins *Eudyptes chrysolophus* (Crossin et al., 2012); tree swallows (Bonier et al., 2011); mourning doves (Miller et al., 2009). Across studies, differences in the relationship between CORT and reproductive success may be due to various fitness measures used, the relative importance of paternal investment, or environmental variation.

While female tree swallows are solely responsible for egg incubation, nestling provisioning is shared with the male (Winkler et al., 2011). As a result, variation in paternal quality may obscure relationships between maternal CORT and investment during the nestling stage. A lack of relationship between CORTnest and female nest box visits has been found in bluebirds (Davis and Guinan, 2014) and other populations of tree swallows (Patterson et al., 2011), suggesting variation among females in their glucocorticoid levels may not directly reflect maternal behaviour. In contrast, Madliger and Love (2016b) found that higher baseline CORTnest in female tree swallows correlated with lower rates of maternal provisioning; however, males compensated for the females’ low rates by increasing their own provisioning rates such that nestlings were not affected. Similarly, Patterson et al. (2011) suggested male tree swallows could compensate for decreased provisioning of their mates, although no male compensation for reduced maternal performance has also been found (Hogle and Burness, 2014). Given the importance of male provisioning to nestling mass gain (Lendvai and Chastel, 2010; Madliger and Love, 2016b; Nomi, 2018), male nest box visits and paternal quality need to be considered when predicting a pairs’ reproductive investment in a nest. Future studies should include male CORT levels, and their relationship with male feeding rates and reproductive success, as in (Ouyang et al., 2011).

The directionality of the relationship between maternal CORT and fitness varies among life stages, populations and species (e.g., Bonier et al., 2009a). Some of this variation is presumably due to the context-dependency of the CORT-fitness relationship and variation in environmental conditions (Burtka et al., 2016; Madliger and Love, 2016b). For example, experimental elevation of maternal CORT levels increased brood mortality in tree swallows, but only when weather conditions were benign (Ouyang et al., 2015). Our inability to detect relationships between maternal hormone levels during chick rearing and reproductive success could be due to the influence of such factors as food availability or weather, both of which could affect body condition and reproductive success of the mother (Schoech et al., 2007; Madliger and Love, 2016a). Maternal baseline CORT may also depend on the habitat type in which female tree swallows were breeding (Madliger and Love, 2016b). While we found no significant difference in reproductive investment between the two study sites, we did find that CORTinc and fledging success tended to be higher at one of our sites (Sewage Lagoon). Reproductive success may perhaps be mediated by a relationship between CORT and foraging conditions (Henderson et al., 2017), which could change from incubation to the nestling stage.

### No relationship between corticosterone and return rates

We found a borderline (p = 0.054) positive relationship between CORTinc and the probability of whether a female returned to the breeding sites the following year. One explanation for the lack of significance is that the relationship between CORT levels and return rates may be non-linear. For example, in cliff swallows (*Petrochelidon pyrrhonata*), highest return rates were seen in individuals with intermediate baseline CORT levels, which could be due to stabilizing selection on CORT levels acting against the detrimental effects of very high or low CORT (Brown et al., 2005; Bonier et al., 2009a). Additionally, environmental variables (Clark et al., 2018) or an individual’s reproductive success may better predict return rates than baseline CORT: the positive (albeit non-significant) effect of fledgling number on maternal return rates that we detected suggests that females with higher reproductive success are more likely to return to a certain area to breed (Bonier et al., 2009b). Thus, CORT may affect return rates and survival indirectly, by affecting fledging success (Shitikov et al., 2017; Weegman et al., 2017).

### Efficacy of silastic implants to raise plasma glucocorticoids

We implanted pre-egg laying females with corticosterone-filled silastic implants, but when females were recaptured during early-to mid-incubation (mean ± SE: 17.0 days ± 0.6 after implantation), the baseline CORT levels of implanted birds did not differ from unmanipulated birds. However, CORT-implanted individuals had a lower probability of recapture during incubation, consistent with a negative relationship between experimental CORT elevation and survival (Schoenle et al., 2019 preprint). Across species, silastic implants have been successfully used to raise CORT levels for anywhere from a few days (Astheimer et al., 2000; Hayward and Wingfield, 2004; Criscuolo et al., 2005; Martin et al., 2005; Angelier et al., 2007) to three weeks post-implantation *in vivo* (Ouyang et al., 2013) and *in vitro* (Newman et al., 2010). However, the use of implants to raise CORT levels has not been consistently successful (Crossin et al., 2012; Ouyang et al., 2013; Hau and Goymann, 2015; Lattin et al., 2016, Torres-Medina et al., 2018). Although the implants used in our study may have failed to release CORT, this seems unlikely given that *in vitro* studies have shown that CORT continues to be released across the membrane over 4 weeks (Newman et al., 2010). More likely, the implants resulted in decreased secretion of endogenous CORT via negative feedback, or increased clearance of CORT from the blood via increased excretory activity (Newman et al., 2010; Henriksen et al., 2011; Robertson et al., 2015).

Rather than experimentally manipulate CORT levels via implants, an alternative approach may be to manipulate maternal condition, such as with feather clipping (Rivers et al., 2017), predator experiments (Clinchy et al., 2011; Pitk et al., 2012), or density manipulations (Bentz et al., 2013).Such an approach would encompass how maternal CORT levels change based on how each female perceives her condition/ environment, how that is reflected in blood CORT levels, and how those levels might influence the next generation (Madliger and Love, 2016a; Rivers et al., 2017).

### Conclusions

The differing directionality of relationship between CORT and fitness among studies and species raises the simple question: is there is a consistent relationship to be found among individuals within a population? Many factors can affect both CORT and fitness, including condition (Love et al., 2005), life-history stage (Romero, 2002), weather (Pakkala et al., 2016), habitat variability (Madliger and Love, 2016b), and resource availability (Breuner and Burk 2019). If it can be reasonably assumed that these will always differ among individuals, then perhaps there is no consistent relationship, and any that may be detected will always be context-dependent (Madliger and Love, 2016a). Recent meta-analyses that seek to understand relationships between CORT and fitness across taxa, and studies that identify factors that contribute to context-dependence, are particularly valuable (Sorenson et al., 2017; Breuner and Burk 2019; Schoenle et al., 2019 preprint; Bonier & Cox 2020).

The use of integrative measures of CORT may be an alternative way to improve our understanding of the relationship between CORT and fitness. By measuring CORT deposited in feathers during growth, or metabolites excreted in feces, it may be possible to infer CORT levels over multiple days of the incubation or nestling stage (Lucas et al., 2006; Bortolotti et al., 2008; Romero and Fairhurst, 2016). For example, giant petrels that successfully bred had higher feather CORT levels than failed breeders, but were less likely to breed the following year, a pattern which was not observed using plasma CORT from these same individuals (Crossin et al., 2013). Ideally, studies could be extended over the winter, as has been done recently in adult tree swallows (Vitousek et al., 2018). This would help elucidate the longer-term effects of maternal CORT on offspring and maternal and fitness.

## ACKNOWLEDGEMENTS

We thank Noah Ben-Ezra, Chantelle Penney and Aleesa Manax for their many hours of help in the field. Erica Nol, Joe Nocera, Jeff Bowman and an anonymous referee provide numerous for suggestions for improving the clarity of an earlier draft of the manuscript.

