## Supplemental Table 1 for "Maternal glucocorticoid levels during incubation predict breeding success, but not reproductive investment, in a free-ranging bird"

**ESM: Table S1. Global statistical models describing variation in female tree swallows in each treatment group (Sham, CORT, and Control). Statistical and mean values (raw mean ± SE) are from results of linear models. Females allocated to each treatment did not differ statistically from each other.**

| Response variable | Sham | CORT | Control | Fixed effects | β | SE | df | t | P | R^2^ |
| --- | --- | --- | --- | --- | --- | --- | --- | --- | --- | --- |
| Pre-laying body mass (g) | 21.3±0.43  N=19 | 22.1±0.46  N=13 | --- | Intercept  Treatment (Sham)  Age (ASY)  Site (Nature Area)  Year | 23.093  -0.783  -0.411  0.661  -0.903 | 0.683  0.593  0.665  0.585  0.572 | 1, 27  1, 27  1, 27  1, 27  1, 27 | 33.811  -1.321  -0.618  1.129  -1.578 | <0.001  0.198  0.542  0.269  0.126 | 0.101 |
| Flattened wing length (mm) | 114.9±0.66  N=20 | 115.7±0.72  N=14 | 116.6±0.71  N=15 | Intercept  Treatment (CORT)  Treatment (Sham)  Age (ASY)  Site (Nature Area)  Year | 116.806  -0.980  -1.699  2.022  0.020  -2.334 | 0.963  1.042  1.022  0.870  0.781  0.762 | 1, 43  1, 43  1, 43  1, 43  1, 43  1, 43 | 121.327  -0.940  -1.662  2.323  0.026  -3.062 | <0.001  0.352  0.104  **0.025**  0.979  **0.004** | 0.169 |
| Clutch initiation date  (Julian day) | 142.9±1.19  N=20 | 142.9±1.32  N=14 | 141.3±1.13  N=20 | Intercept  Treatment (CORT)  Treatment (Sham)  Age (ASY)  Site (Nature Area)  Year | 144.917  1.581  1.583  -7.027  2.679  2.458 | 1.704  1.768  1.697  1.480  1.387  1.361 | 1, 48  1, 48  1, 48  1, 48  1, 48  1, 48 | 85.059  0.894  0.933  -4.747  1.931  1.806 | <0.001  0.376  0.356  **<0.001**  **0.059**  0.077 | 0.339 |

Control birds did not have a pre-laying body mass measured because they were not captured until incubation

Significant main effects are bolded.
